# Calredoxin regulates the chloroplast NADPH-dependent thioredoxin reductase in *Chlamydomonas reinhardtii*

**DOI:** 10.1101/2022.11.22.517551

**Authors:** Karen Zinzius, Giulia Maria Marchetti, Ronja Fischer, Yuval Milrad, Anne Oltmanns, Simon Kelterborn, Iftach Yacoby, Peter Hegemann, Martin Scholz, Michael Hippler

## Abstract

Calredoxin (CRX) is a calcium (Ca^2+^)-dependent thioredoxin (TRX) in the chloroplast of *Chlamydomonas reinhardtii* with largely unclear physiological role. We elucidated the CRX functionality by performing in-depth quantitative proteomics of wild type cells in comparison with *crx* insertional mutant (IM_*crx*_), two CRISPR/Cas9 KO mutants and CRX rescues. These analyses revealed that the chloroplast NADPH-dependent TRX reductase (NTRC) is co-regulated with CRX. Electron transfer measurements revealed that CRX inhibits NADPH-dependent reduction of oxidized chloroplast 2-Cys peroxiredoxin (PRX1) via NTRC and that the function of the NADPH-NTRC complex is under strict control of CRX. Via non-reducing SDS-PAGE assays and mass spectrometry, our data also demonstrated that PRX1 is more oxidized under high light (HL) conditions in the absence of CRX. The redox tuning of PRX1 and control of the NADPH-NTRC complex via CRX interconnects redox control with active photosynthetic electron transport and metabolism as well as Ca^2+^ signaling. In this way, an economic use of NADPH for PRX1 reduction is ensured. The finding, that the absence of CRX under HL conditions severely inhibited light-driven CO_2_ fixation underpins the importance of CRX for redox tuning as well as for efficient photosynthesis.

**One-sentence summary:** Calredoxin dependent redox regulation ensures efficient photosynthesis.

## Introduction

The process of oxygenic photosynthesis provides an important basis for life on earth, because it drives the conversion of light energy into chemical energy that is used for building up complex organic material.

The first steps of plant photosynthesis mediate the photoinduced electron transfer from water to NADP+, catalyzed by photosystems I and II (PSI and PSII) of the photosynthetic electron transfer chain. PSI and PSII are interconnected by the cytochrome *b_6_/f* complex, which transfers the electrons from PSII to PSI and translocates protons into the thylakoid lumen. This photosynthetic process, also termed ‘light reactions’, drives light-dependent water oxidation, NADP^+^ reduction and ATP formation (Whatley, et al., 1963). Formation of ATP is catalyzed by the ATP synthase driven by the proton-motive force generated by the light reactions (Mitchell, 1961). ATP and NADPH are used by the light-independent ‘dark reactions’ of the Calvin-Benson-Bassham cycle (Bassham, et al., 1950) to assimilate CO_2_. During the evolution photosynthesis was optimized to increase photosynthetic performance but at the same time to minimize photo-oxidative stress and to avoid an excess formation of reactive oxygen species (ROS). Several signaling mechanisms, including calcium (Ca^2+^) and redox signaling balance the absorption of light energy with downstream energy consuming pathways such as CO_2_ fixation and conversion (Foyer, 2018; Hochmal, et al., 2015). Once ROS are generated, several proteins relay this information to target proteins via redox signaling. At the same time, these proteins reduce and thereby detoxify ROS, thus commonly referred to as ‘ROS detoxifying’ or ‘ROS processing enzymes’ (Noctor, et al., 2018). One important class of H_2_O_2_ interacting proteins within the chloroplast are peroxiredoxins (PRXs) (Dietz, 2003; Poole et al., 2004; Dietz et al., 2006). These proteins harbor varying numbers of cysteines forming intramolecular disulfide bridges when oxidized by ROS. Interestingly, it has been recently discovered that PRXs, alongside their “antioxidative” action, can also act as sensor relays, transferring the oxidizing equivalents from H_2_O_2_ to their interacting partners favouring the transmission of the oxidative stress response (Sobotta et al., 2015), and, third, are also involved in the oxidation of chloroplast enzymes in the dark (Ojeda et al., 2018). When oxidized, regeneration of PRXs is accomplished by reduction via the thioredoxin (TRX) system (Pulido et al., 2010; Pascual et al., 2011) (Dietz, 2011). TRXs use redox active cysteines for target reduction while regaining their electrons from either reduced ferredoxin (Fd) via ferredoxin-TRX-reductases (FTRs) or NADPH via NADPH-dependent TRX reductases (NTRs). In the chloroplast of algae, plants as well as in some cyanobacteria, an atypical NTR with a TRX domain fused to its C-terminus can be found (Serrato, et al., 2004). This enzyme, termed NTRC, functions as a homodimer (Perez-Ruiz and Cejudo, 2009) and is also able to directly reduce PRXs efficiently. Beside its antioxidant function (Perez-Ruiz et al., 2006), which has been shown to be of vital importance in *A. thaliana* especially under low irradiance (Carrillo et al., 2016), NTRC is involved in the redox regulation of a plethora of different metabolic pathways. These include the CBB cycle (Thormahlen et al., 2015), chlorophyll (Richter et al., 2013) and starch biosynthesis (Lepisto et al., 2013) and even fruit growth (Hou et al., 2019). The crystal structure of the *Chlamydomonas reinhardtii* (Chlamydomonas) NTR domain of NTRC was solved only recently (Marchetti et al., 2022).

Furthermore, Calredoxin (CRX), a chloroplast-localized TRX-like protein interacting with the 2-cysteine peroxiredoxin PRX1, was found to be involved in redox regulation and ROS scavenging, after being induced by autotrophic and/or high light (HL) conditions (Hochmal, et al., 2016). CRX is a combination of a TRX with a Ca^2+^-binding domain, the latter controling the redox activity of the TRX domain *in vitro*. Decrease of CRX amounts in an insertional mutant (IM_*crx*_) led to increased lipid peroxidation and cyclic electron flow around PSI *in vivo* (Hochmal, et al., 2016). As the redox potential of PRX1 is slightly more negative (−310 mV (Yoshida and Hisabori, 2017)) than the redox potential of CRX (−290 mV (Hochmal et al., 2016)), reduction would be on the side of CRX, which by itself would drive partial oxidation of PRX1. However, electron transfer measured *in vitro* between CRX and PRX1 showed efficient reduction of PRX1 probably due to the negative potential of NADPH and the fast consumption via H_2_O_2_ reduction (Hochmal et al. 2016). Electron exchange between CRX and PRX1 in the one or other direction should therefore rely on the individual redox and Ca^2+^ status of the microenvironment. Interestingly, it was shown that *in vitro* electron transfer to PRX1 is similar for CRX and NTRC in the presence of Ca^2+^ (Marchetti et al., 2022). Notably, the NTRC midpoint redox potential was found at −275 mV (Yoshida and Hisabori, 2016), which is slightly lower as compared to the one of CRX.

To gain further insights into CRX function, we performed in-depth quantitative proteomics analyses taking advantage of the *crx* insertional mutant (IM_*crx*_), two CRISPR-Cas9-generated knockout mutants (*crx* KOs, E1 and A5) and two complemented strains (IM-R and A5R). The data revealed a co-regulation of CRX and NTRC, underpinning involvement of CRX in the chloroplast redox homeostasis of *Chlamydomonas*. The absolute quantitation of these two proteins and their use in *in vitro* redox assays interestingly highlighted an inhibitory role of CRX towards the general redox activity of NTRC and in particular towards its ability to reduce 2-Cys PRX1 *in vivo*. Taken together, our results highlight the possible role of CRX as a master regulator in the highlight stress response in the chloroplast by regulating the expression and the activity of NTRC.

## Results

### Generation of *crx* KO mutants via CRISPR/Cas9 and phenotypic analyses

To reveal new insight into the function of CRX, two *crx* KO strains were generated from WT CC125 via CRISPR-Cas9 and were back-crossed three times with the parental strains CC124 (mt−) and CC125 (mt+) (Figure 1). Importantly, these KO strains (E1 and A5) lacked residual CRX accumulation, unlike the previously described insertional CRX mutant IM_crx_, which showed about 10-20 % CRX expression compared to the WT (Hochmal et al., 2016). The back-crossed A5 strain as well as the IM_crx_ were complemented by transformation with a vector harboring genomic DNA from 900 bp upstream until 300 bp downstream of the coding region in order to include regulatory motifs, such as the promotor region for *crx*. The complemented strains were selected for maximum accumulation of CRX by protein immunoblotting. For each *crx* mutant, one complemented strain having more than 50% of CRX expression as compared to WT was selected for further experimental analysis (A5-R and IM-R).

**Figure 1:**
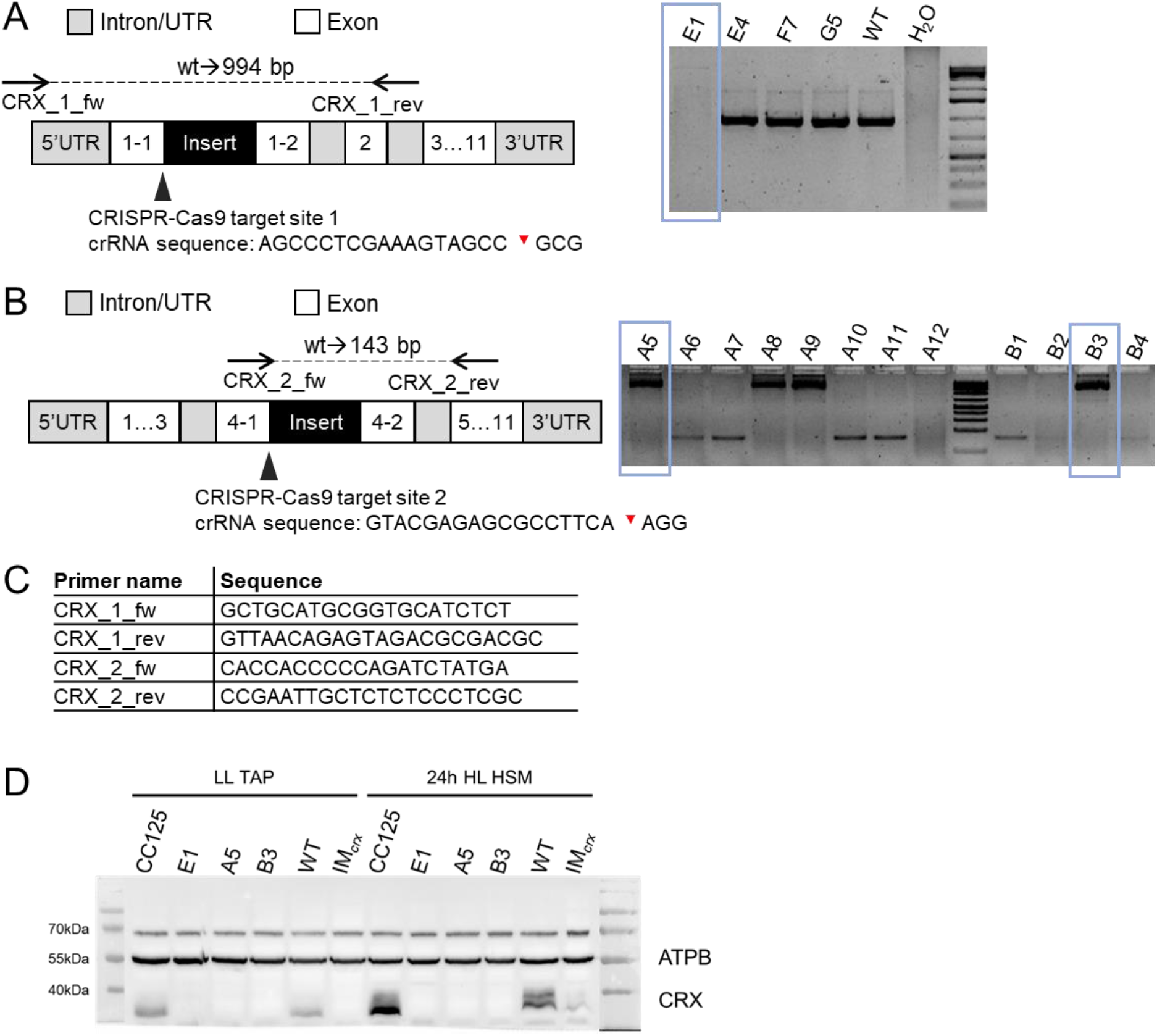
CRISPR-Cas9 mediated KO of CRX. Generation of CRX mutant affected in exon 1 (A). Only the clone, E1, showed a PCR product bigger in size than WT. A second set of mutants was created by targeting a sequence in exon 4 (B). Sequencing of PCR products deviating from WT product size showed that all respective mutants are isogenic, having part of the co-transformed paromomycin vector integrated after the predicted Cas9 cleavage site. Primer sequences used for mutant screening (C). For final proof of knockout, a chosen set of mutant and WT strains were subjected to a TAP/LL to HSM/HL (~200 μEm^−2^s^−1^) shift. Cells were harvested, lysed and after separation by SDS-PAGE, proteins were transferred onto a nitrocellulose membrane and probed with antibodies against ATPB and CRX (D). As mutants A5 and B3 turned out to be isogenic while both are deficient in CRX, only mutant A5 was used for further experiments. Cas9 cleavage sites within the crRNA sequence are marked by red triangles in A and B.

When examining the phenotypes, the first thing that became apparent was that wildtype, mutant and complemented strains differed considerably in their ability to grow at high light (HL) intensities under photoautotrophic growth conditions (Figure 2). Different dilutions of the WT and backcrossed mutant and complemented strain cultures spotted onto agar plates showed comparable growth in low light (LL), but an impaired growth of the mutants in comparison to the respective wild types in HL. This effect is strongest for E1, followed by A5 whereas IM_crx_ is least affected. The complemented strains showed an intermediate growth phenotype between respective *crx* mutants and WTs. The growth of WT CC125 was also affected in TP high light but not under HSM high light growth conditions.

**Figure 2:**
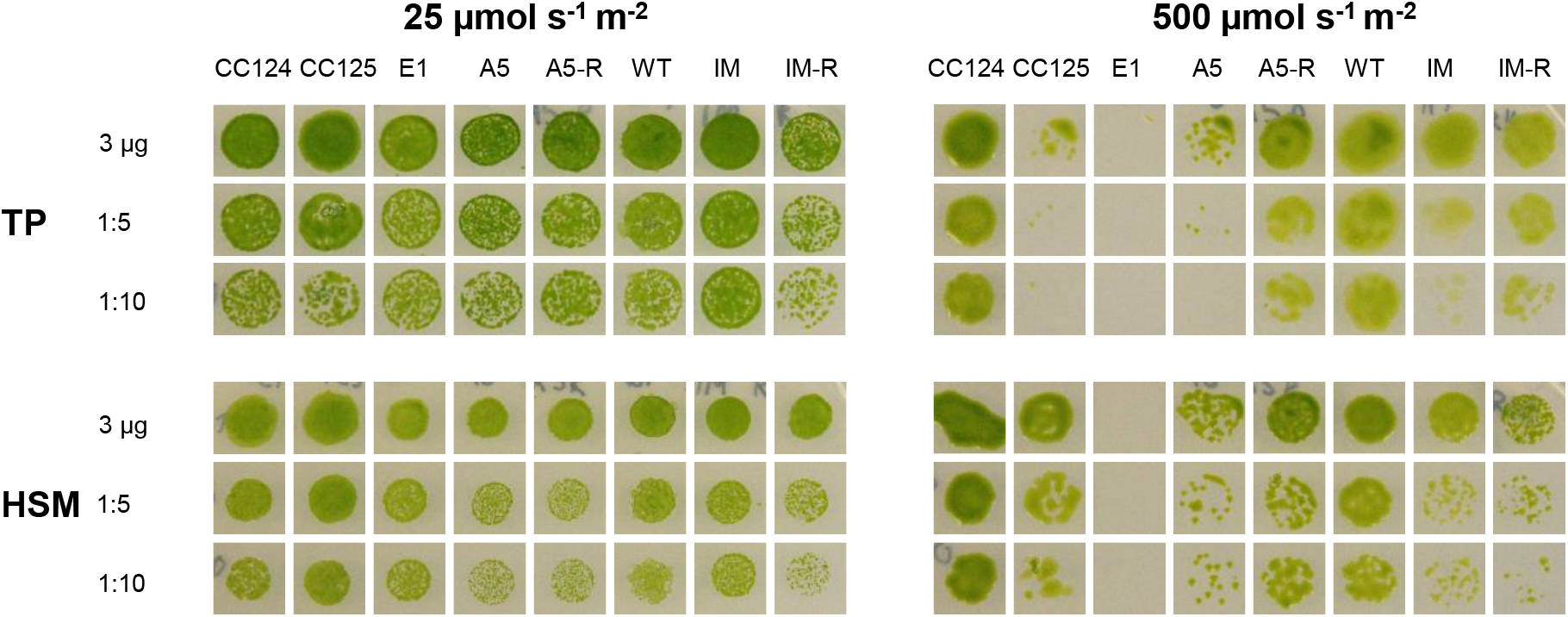
Lack of CRX leads to decreased growth in HL conditions. Growth performance of back-crossed CRISPR-Cas CRX mutants A5 and E1 and the insertional IM_crx_ was compared to the respective parental strains (CC124, CC125 and CC4375 (WT_crx_)) and resuces (A5-R and IM-R) by spotting 10 μl of culture at 3 μg/ml chlorophyll and the corresponding dilutions onto tris-phosphate (TP) or high salt medium (HSM). Pictures were taken after 1 week of incubation at low (25 μmol m^2^ s^−1^) or high (500 μmol m^2^ s^−1^) light.

### Quantitative proteomic analyses to identify proteins co-regulated with CRX

To unravel the underlying mechanisms of the CRX HL phenotype, the proteomes of the backcrossed CRX mutants, corresponding wild types and complemented strains were analysed by mass spectrometry. For each strain, four independent biological replicates were measured. Synchronized cells were grown in TP medium in LL and then shifted to HL. Samples were taken at 0 (LL samples) and 24 h (HL samples) after the switch to HL. Whole protein was extracted from the samples, tryptically digested, and analysed by HPLC-MS/MS. Out of 5861 detected proteins, 881 were identified as differentially regulated in the different strains and submitted to hierarchical clustering (Figure 3A). Among the clusters identified (Supplementary Table 1), those relevant to the phenotype are shown in detail in Figure 3B and C. Several proteins belonging to PSI and PSII as well as ATPase were found to be significantly downregulated in the strain E1 in comparison to the respective WT (CC125) (Figure 3B), whereas these proteins were not affected or even upregulated in A5. Analyses of the growth phenotypes (Figure 2) revealed, that growth was strongly impaired in E1 and A5, but also of CC125 was more affected in HL TP, while in HSM, in contrast to E1, growth of CC125 was not light sensitive. Interestingly, a significant downregulation of PSII and ATPase subunits is also observed in the IM_crx_ strain and rescued in the complemented IM-R strain (Figure 3B). In line, PSII quantum efficiency and O_2_ evolution were severely affected in E1 and IM_crx_ but not A5 and IM-R (Supplementary Figure 1). Remarkably however, CO_2_ fixation in HL, as measured via membrane inlet mass spectrometry (MIMS), was affected not only in E1 but also in A5 and IM_crx_, pointing towards another phenotype, beside the growth defect, that is correlated with the lack of CRX.

**Figure 3:**
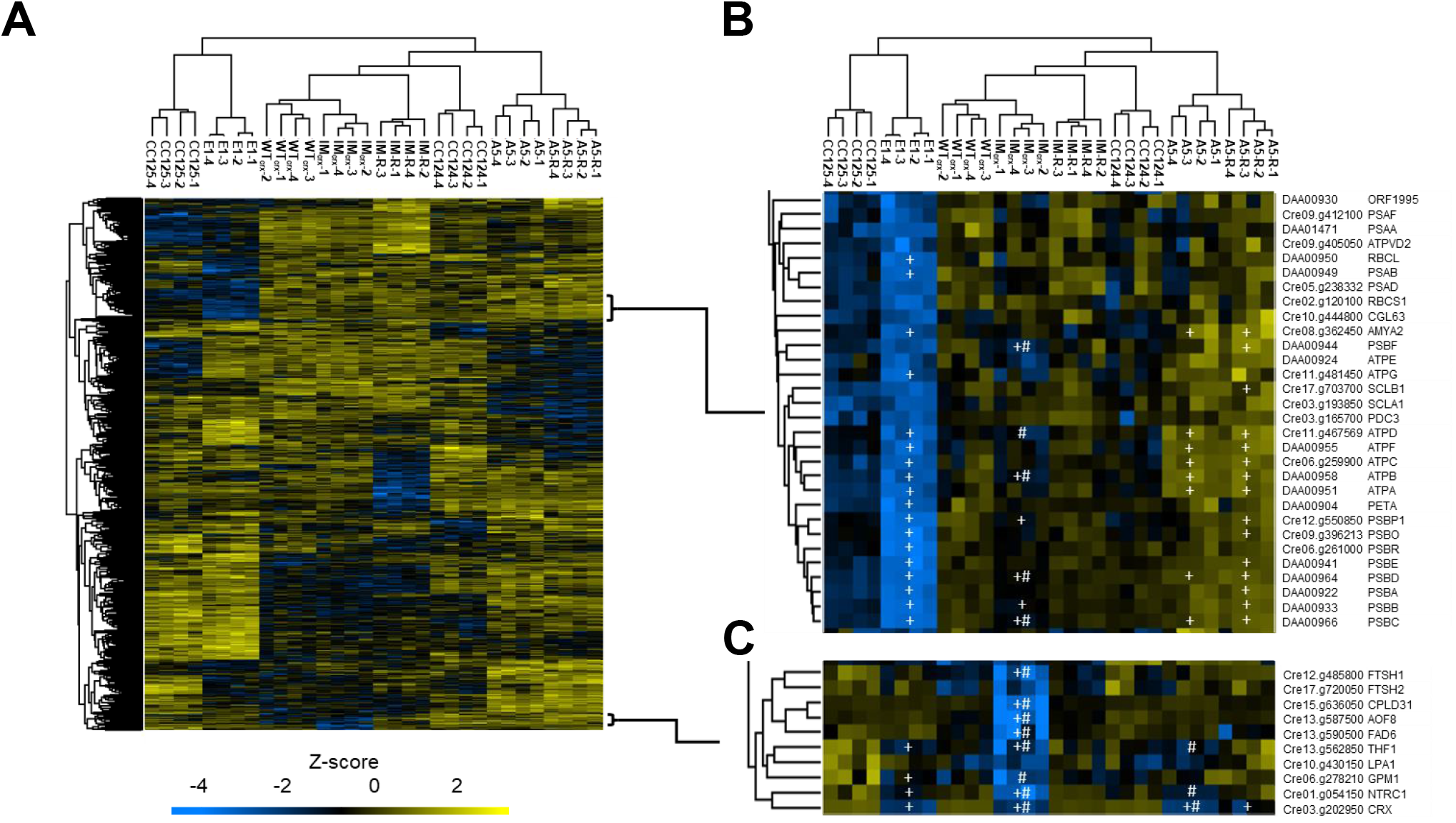
Lack of CRX induces decrease in NTRC accumulation. Hierarchical clustering of 881 proteins with significant differential abundance (p<0.01, ANOVA followed by Tukey’s posthoc test with FDR of 1%) after 24 h high light (500 μmol·m^−2^·s^−1^) (A). Downregulation of co-regulated proteins belonging to PSI, PSII and ATPase in the E1 and CC125 strains (B). Proteins co-clustering with CRX (C). In the highlighted clusters, proteins with significantly different expression between either mutant/rescue vs corresponding wt (+) or mutant vs rescue (#) are indicated.

The other cluster of interest (Figure 3C) contains all proteins co-regulated with CRX. c*rx* was significantly less expressed in all mutants, including IM_crx_, while the protein was significantly more abundant in the complemented strains, particularly in IM-R. The only partial re-establishment of *crx* expression in A5-R is in agreement with the spot test, where the growth performance of this strain was only partially restored. Interestingly, another protein involved in photosynthetic redox regulation belongs to this cluster, namely NTRC (Cre01.g054150). This protein has a pattern of expression highly similar to the one of CRX, being significantly less expressed in the *crx* mutants but re-established in the CRX complemented strains. The other protein which showed a similar behaviour is thylakoid formation 1 (THF1) (Cre13.g562850).

In conclusion, among the 881 differentially expressed proteins, given the distinct genetic backgrounds of E1, A5 and IM_*crx*_, only NTRC and THF1 expression patterns were found to correlate with CRX expression. This strongly suggests a functional link between these proteins and CRX. As NTRC is a hub of chloroplast redox regulation in vascular plants and a potential redox partner of CRX, we focused on the putative functional interconnection between these two proteins. The significant downregulation of NTRC in the *crx* mutants was also observed when quantitative proteomics data of all mutants are compared to all WTs via independent Vulcano plot analyses (Supplementary Figure 2A). It is also of note that the comparison of CC124 and CC125, as revealed by Vulcano plot analyses, showed significant differences in protein expression, e.g. enzymes of acetate metabolism and a glutathione peroxidase were more abundant in CC124, while e.g. VIPP1 as well as ALD5 were more abundant in CC125 (Supplementary Figure 2B).

### Redox activity measurements of electron transfer between CRX, NTRC and PRX1

As both proteins are known to function as oxidoreductases, electron transfer between the two proteins was analysed in redox activity measurements (Figure 4). Interestingly, the NTRC reduction capacity is negatively affected by CRX, as DTNB (Ellman’s reagent) or PRX1 reduction is decreased with increasing concentrations of CRX (Figure 4A+D). This inhibition is independent of the CRX redox state, as also the redox inactive CRX mutant C1,2S led to slower DTNB reduction. Furthermore, the lack of Ca^2+^ releases the inhibition partially (Figure 4A), making Ca^2+^ an important compound for the NTRC redox inhibition. The inhibition is specific to CRX, as shown by the fact that another redox active protein, TRXf, had no negative impact on NTRC activity (Figure 4B). On the contrary, the CRX reduction capacity was unaffected by the presence of NTRC-C136S, a mutated version with redox-inactive NTR domain, but active TRX domain, when electrons were donated by *E. coli* TRX reductase (Figure 4C).

**Figure 4:**
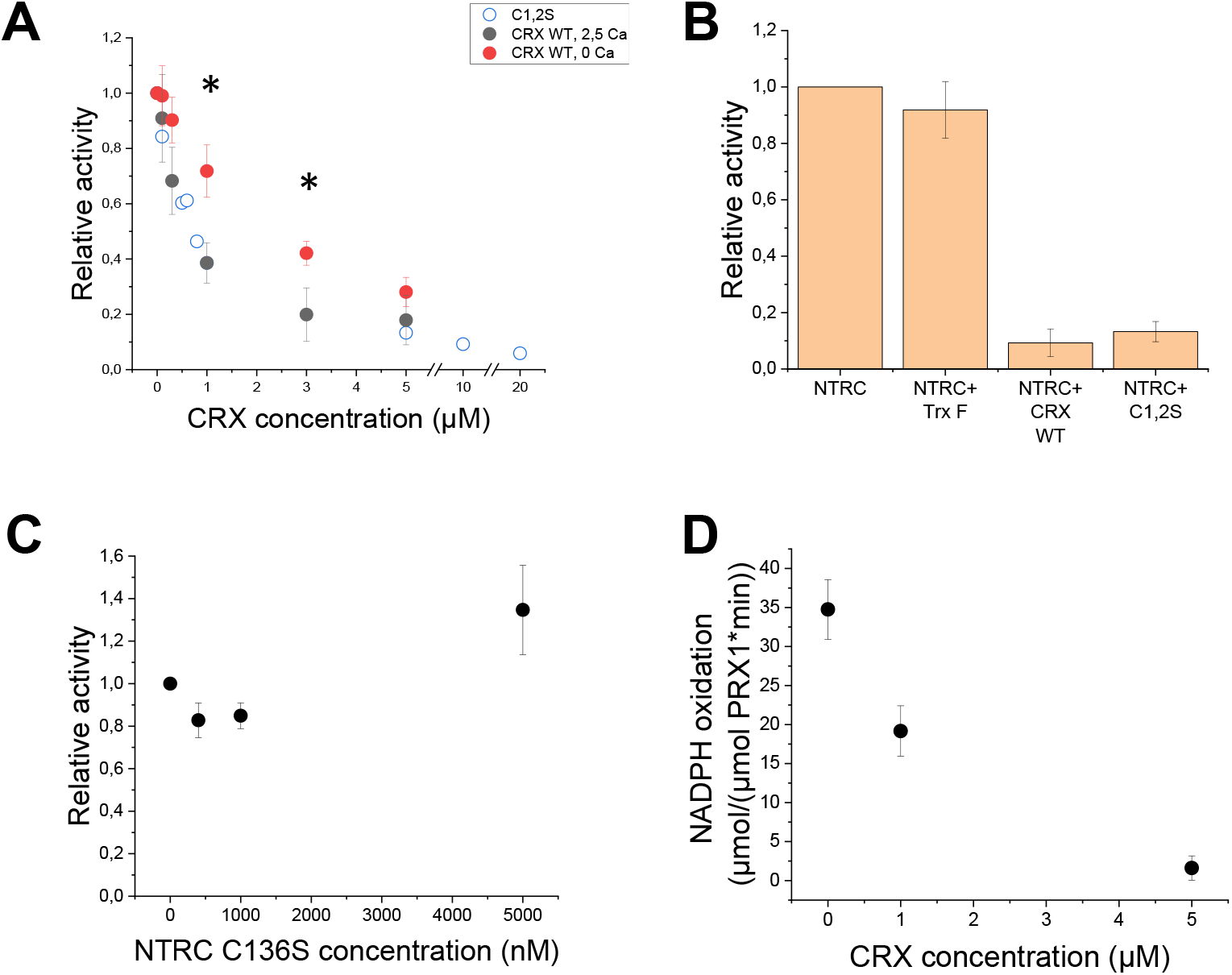
CRX specifically inhibits NTRC redox activity. 400 nM recombinant NTRC were incubated for 10 minutes at RT in presence of either 0 (black dots) or 2.5 μM (red and empty dots) free calcium, 200 μM NADPH, and different concentrations of either WT CRX (full dots) or its active site mutant (C1,2S, empty dots) (A). After ten minutes 200 μM DTNB were added as substrate for reduction. The increase in absorption at 412 nm was recorded to calculate the redox activity (slope 0-80 s after addition of DTNB). Data were normalized on the highest activity measured for each protein purification. Error bars represent s.d. of three independent measurements. Stars indicate a significant difference by student’s t-test of p<0.05. The same settings were used for incubation of 400 nM NTRC and 5 μM of either TrxF, CRX WT or its active site mutant (C1,2S) (B), and for incubation of 1 μM CRX, 200 nM TrxR and different concentrations of NTRC NTR domain active site mutant C136S (C). The reduction of 1 μM oxidized recombinant PRX1 was monitored by measuring NADPH oxidation at 340 nm in presence of 5 μM recombinant NTRC and different concentrations of CRX WT at a concentration of 2.5 μM free calcium in presence of 80 μM H_2_O_2_ (D). Error bars represent s.d. of three independent measurements.

### Absolute quantitation of CRX and NTRC

To relate these functional properties of CRX and NTRC to a putative *in vivo* situation, absolute protein quantification was performed using protein samples obtained from algae cultivated in TP medium (Figure 5). To do so, 50 ug whole cell extract of TP grown CRX WT or mutant cells were tryptically digested with 25, 125 or 500 fmol recombinant ^15^N-labelled CRX and NTRC. The different isotope masses allow to distinguish between recombinant and endogenous CRX or NTRC and to relate the known ^15^N protein amounts to the unknown ^14^N protein amount from the cell lysate. As previously demonstrated by (Hochmal et al., 2016), CRX expression was enhanced in the wild type strains under high irradiance. Furthermore, no peptides corresponding to CRX were found in E1 and A5, confirming a successful mutation via CRISPR-Cas9 in these strains. In the WTs, CRX accumulation was much higher as compared to NTRC, with an overall ratio of approximately 9 under both low and high light conditions. Furthermore, NTRC expression in the CRX mutants was significantly reduced in high light compared to the WTs, which is in line with results of the relative quantification (Figure 3). Interestingly, this reduction was not present under low irradiance, where NTRC abundance was similar in all strains.

**Figure 5:**
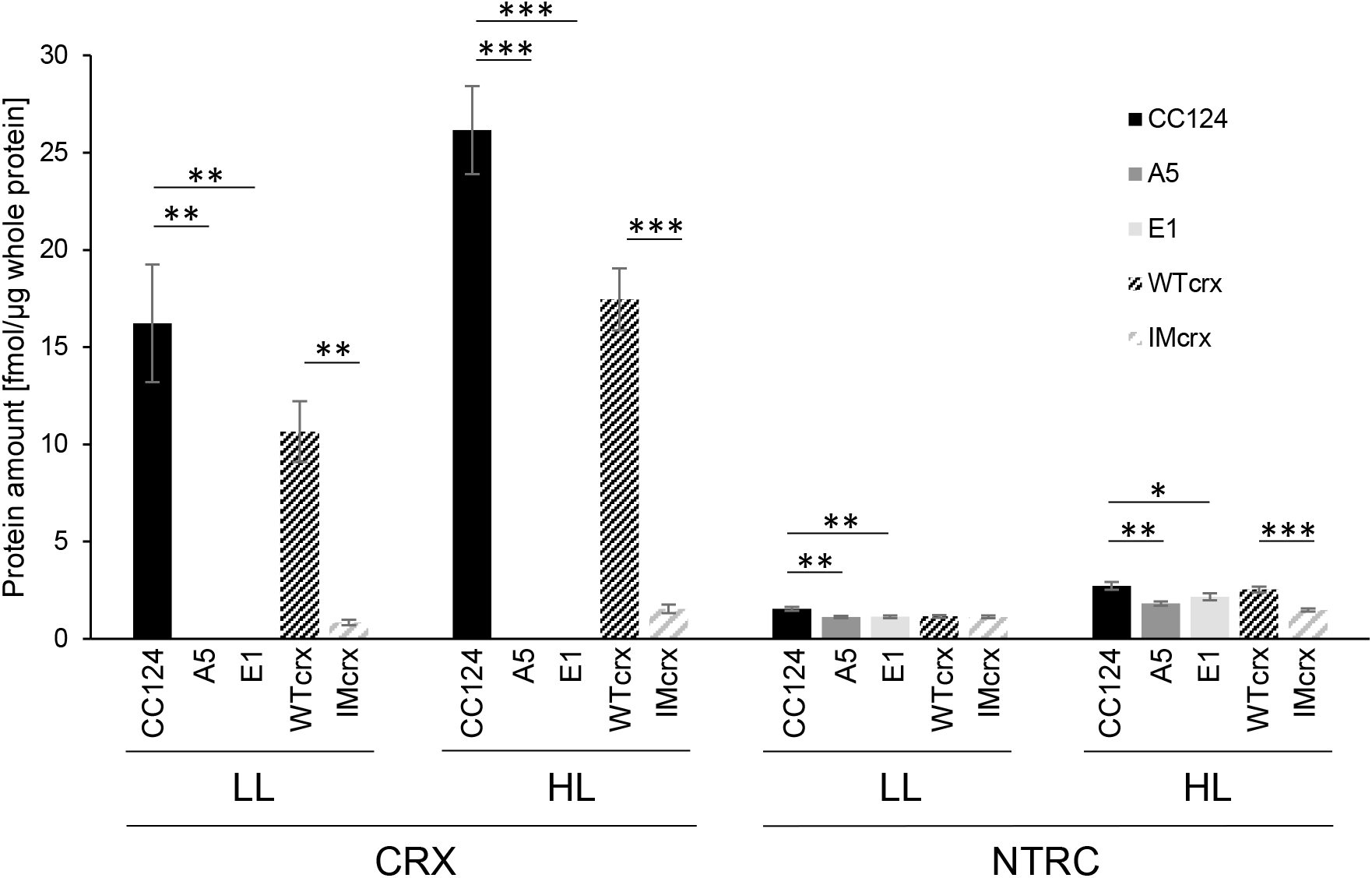
Determination of absolute CRX and NTRC amounts in the wild type strains CC125 and WTcrx (CC4375) in LL (30 μmol·m^−2^·s^−1^) and HL (500 μmol·m^−2^·s^−1^). For absolute quantitation, recombinant ^15^N-labeled CRX and NTRC were spiked into ^14^N whole cell extracts grown in TP medium. Quantitation of intrinsic CRX and NTRC were based on known amounts of ^15^N-labeled CRX and NTRC. Bars indicate s.d. of 3 replicates with different amounts of ^15^N CRX/NTRC added. Stars indicates statistical significance by Student’s T-Test of *<0.05, **<0.005, ***<0.001.

### Functional analyses of CRX and NTRC interactions

To analyze the NTRC-CRX interaction and its impact on the PRX1 redox state, non-reducing SDS-PAGE was performed after incubating the three proteins in different combinations and under varying conditions (Figure 6). Because PRX1 is known to form dimers upon oxidation, the presence or absence of either monomer (~25 kDa) or dimer (~55 kDa) allows visualization of the PRX1 redox state. As long as CRX was absent, NTRC affected the PRX1 redox state as expected (Figure 6A+B): In experiments, where reduced PRX1 was used as substrate, PRX1 stayed reduced, independent of the NTRC redox state and the presence of NADPH (Figure 6B). When oxidized PRX1 was used as substrate, it remained oxidized as long as WT NTRC and NADPH were added, which led to PRX1-reduction using electrons from NADPH (Figure 6A). However, it could be reduced without NADPH when WT-NTRC or NTRC-C136S were already reduced before the experiment. Here, electrons from the reduced TRX domain of WT or NTRC-C136S could directly be transferred to PRX1 (Figure 6A, lane 8-10). In contrast, the NTRC-C136S mutant protein could not reduce PRX1, when NADPH was present (Figure 6A, lane 11, marked with *), which led to the hypothesis that NADPH induces a conformational change making the TRX domain unavailable for PRX1 reduction in NTRC-C136S.

**Figure 6:**
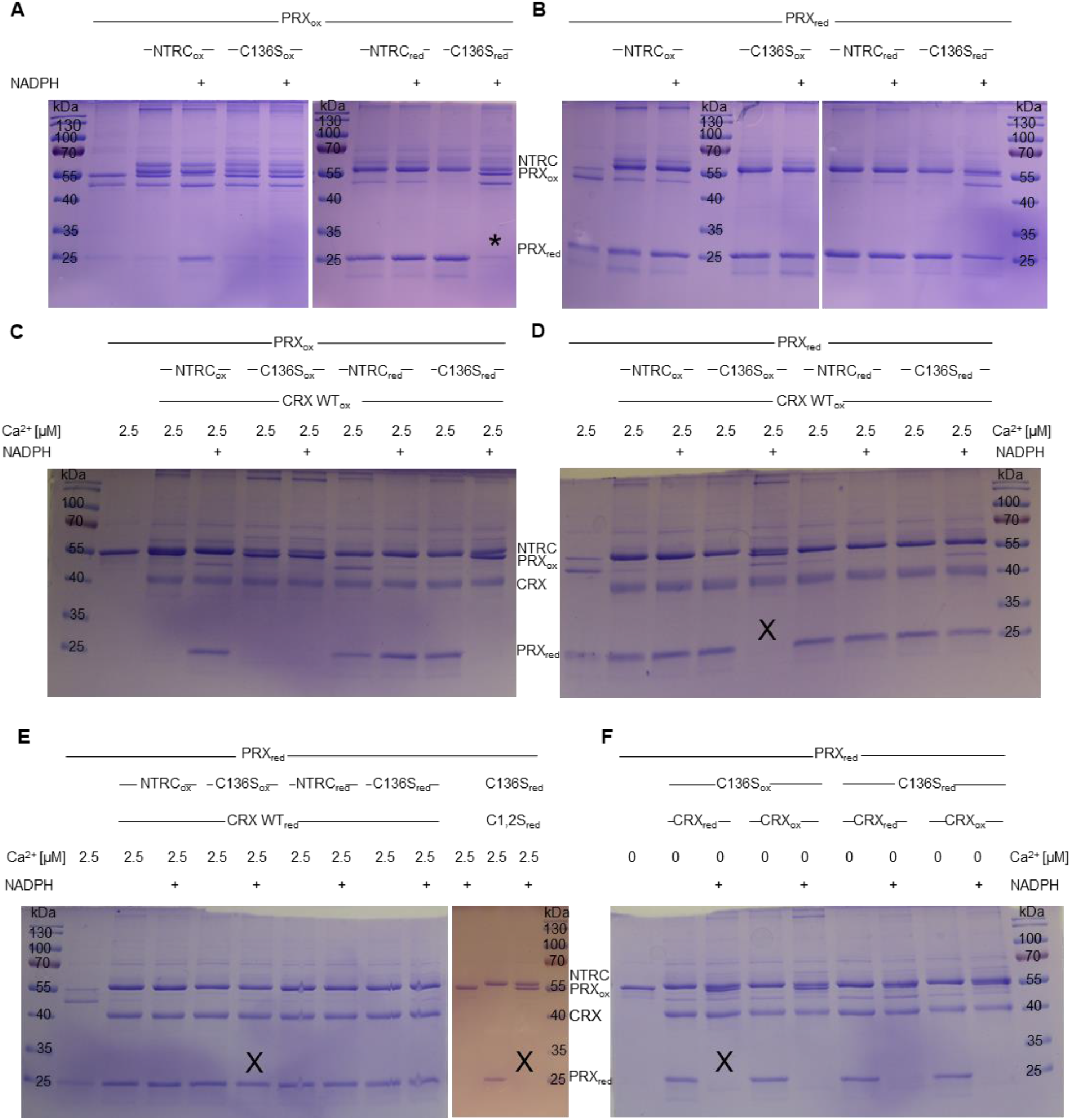
Oxidation of PRX1 is promoted via non-active CRX in the presence of NADPH. 1 μg of purified recombinant PRX1 was reduced (oxidized) with 1 mM DTT (H_2_O_2_) for 1h at room temperature (RT). After filtration over a G25 column, incubation with 1 μg of recombinant, equally reduced (oxidized) and filtered NTRC, NTRC-C136S and/or CRX, CRX-C1+2S for 1h at RT lead to redox exchanges between the different proteins. Presence or absence of Ca^2+^ and/or NADPH (100 μM) during this incubation is indicated. Before loading onto SDS-PAGE gels, addition of non-reducing loading dye and 20 min heating to 70°C ensured denaturation but allowed to distinguish between PRX1 dimers (PRX1_ox_) or monomers (PRX1_red_) after Coomassie staining. Pictures represent results from two independent replicates.

When oxidized CRX was added, the same results were obtained (Figure 6C+D), except for the NTRC-C136S mutant, in the presence of NADPH and previously reduced PRX1 (Figure 6D, lane 5, marked with X). Under these conditions, PRX1 appeared to be oxidized, when oxidized CRX is present, suggesting that in the presence of NADPH either CRX binding makes the oxidized TRX domain of NTRC-C136S available for PRX1 oxidation or that the oxidized CRX itself oxidizes PRX1. However, when reduced CRX was added, this oxidation is probably overcome by re-reduction via CRX (Figure 6E, lane 6, X), as previously reported (Hochmal et al., 2016). As CRX is known to be active only in the presence of Ca^2+^ (Hochmal et al., 2016), addition of EGTA abolished this re-reduction and PRX1 became oxidized even in the presence of reduced CRX (Figure 6F, lane 3, X). Also, the use of reduced CRX double cysteine mutant (C1+2S) did not lead to PRX1 re-reduction (Figure 6E, lane 13, X), underpinning the importance of redox-active CRX for reduction of PRX1. In conclusion, the simultaneous presence of inactive CRX (oxidized, mutated active Cys or absence of Ca^2+^) and NADPH impacted PRX1/NTRC in a way that PRX1 did not stay reduced. Importantly, the lack of Ca^2+^ in the presence of NADPH always led to PRX1 oxidation, independent of the redox state of NTRC-C136S or CRX (Figure 6F). As the NTRC-C136S mutant cannot reduce its own TRX domain, electron replenishment from NADPH is disabled. Therefore, oxidation of PRX1 can be observed instead of being overruled by re-reduction via WT NTRC.As PRX1 is the main CRX/NTRC target, non-reducing gels after NEM labelling were run and analysed for the fraction of reduced PRX1 *in vivo* (Figure 7). Indeed, we observed a significant disability to keep PRX1 in a reduced state in the *crx* KO mutants A5 and E1 in comparison to WT.

**Figure 7:**
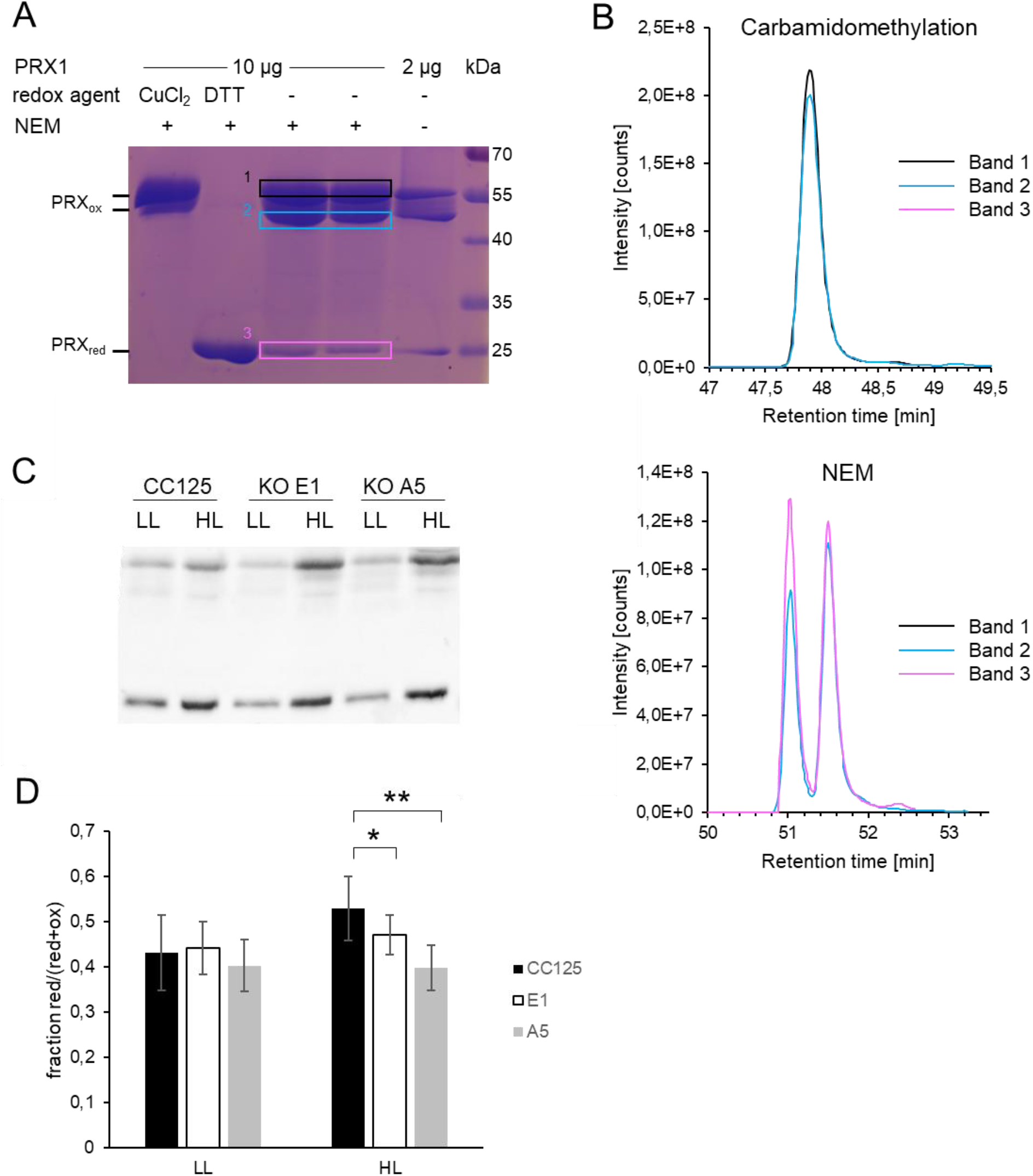
CRX is required to sustain reduction of PRX1 *in vivo*. Recombinant PRX1 was loaded onto a non-reducing SDS-PAGE after optional oxidation (50 μM CuCl_2_) or reduction (100 mM DTT) and NEM labeling (A). The two bands from dimerized PRX1 (bands 1 and 2) and the one representing monomeric PRX1 (band 3) were cut, reduced with DTT, carbamidomethylated and digested with trypsin. Peptides were analyzed by mass spectrometry as described in Material & Methods. Panel (B) shows precursor ion traces of the carbamidomethylated and NEM-labelled PRX1 peptide VLQAIQYVQSNPDEVCPAGWKPGDKTMKPDPK. The two peaks observed for the NEM-labelled peptide most likely represent two diastereomers, which were generated by introducing a new chiral center upon NEM-labelling (Smyth et al., 1964; van den Hooven et al., 2001; Esteban-Fernandez et al., 2011). *In vivo* PRX1 redox state in CRX WT and mutant strains before (LL) and after (HL) a 24 h shift to 500 μEm^−2^s^−1^ (C). Samples were harvested in 10 % TCA to block the thiol redox state and subsequent NEM labeling avoided reoxidation during sample handling. Samples were run on a non-reducing SDS-PAGE and blotted onto a nitrocellulose membrane. PRX1 was detected by a polyclonal antibody raised against the recombinant PRX1. The fraction of reduced PRX1 (intensity reduced band(s) divided by the sum of reduced and oxidized bands within one sample as measured by ImageJ) was calculated and plotted in (D). Stars indicate significant differences by Student’s t-test with a p-value of *<0,05 and **<0,005.

## Discussion

To shed further light onto CRX function, we used two CRX CRISPR/Cas9 knockout mutants and an insertional mutant as well as CRX complemented strains. Analysis of these strains via (relative) quantitative proteomics pointed to a functional link between CRX and NTRC (Figures 3–8). This functional link was further demonstrated by combined enzyme and non-reducing gel assays. Moreover, the data revealed a severe HL growth phenotype in CRX depleted strains (Figures 1+2) as well an impact in CO_2_ fixation under HL conditions (Supplementary Figure 1).

**Figure 8:**
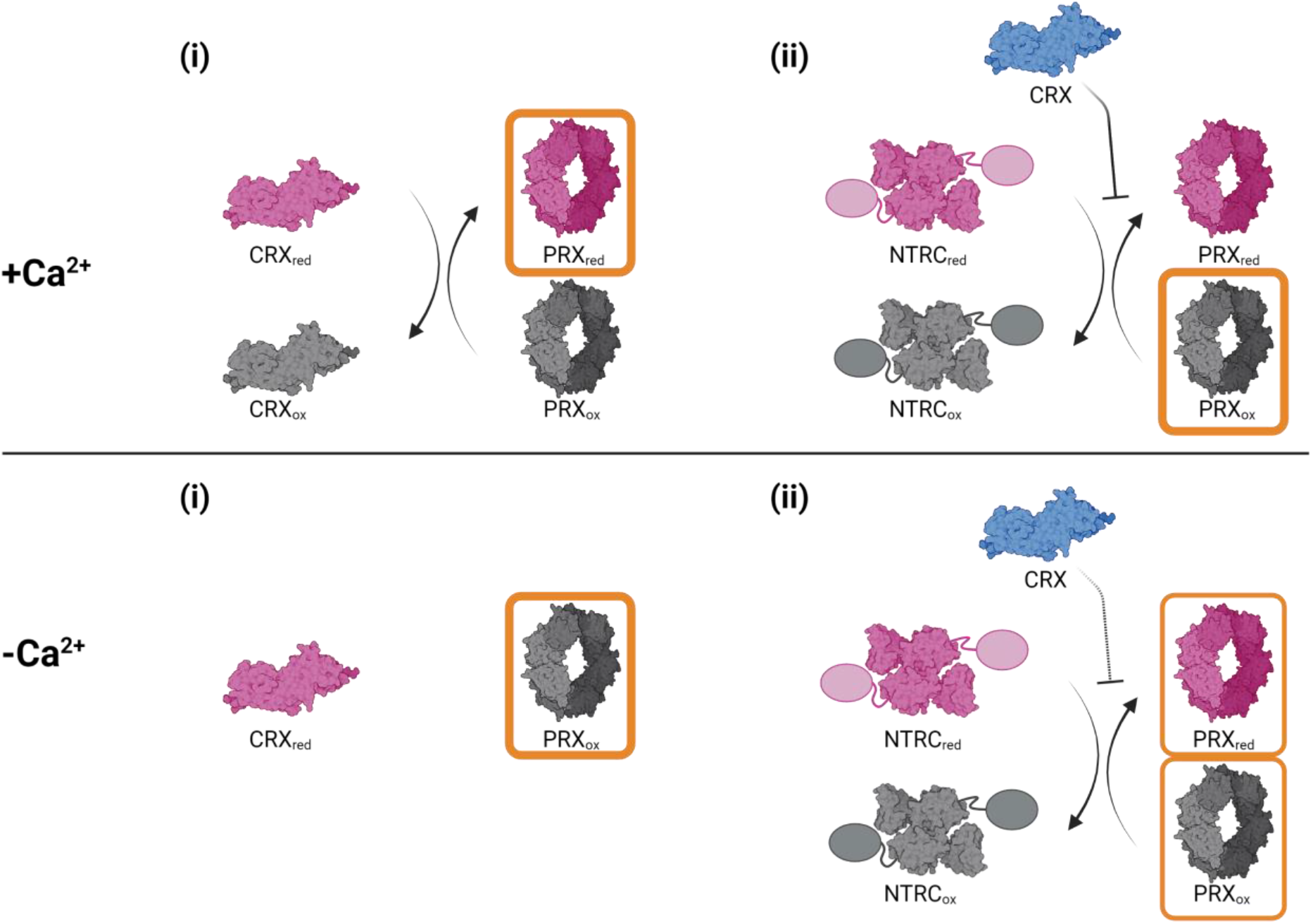
Model for CRX, PRX1 and NTRC interaction in the presence of NADPH. In the presence of Ca^2+^, CRX can reduce PRX1 (i) and also inhibits NTRC reduction of PRX1 (ii). In the absence of Ca^2+^, CRX cannot reduce PRX1 and NTRC inhibition is weaker. The resulting redox state of PRX1 is marked in orange. Figure created with BioRender.com.

### Quantitative proteomics reveals co-regulation of CRX and NTRC

Using mass spectrometry and subsequent hierarchical clustering of protein accumulation profiles, CRX was found to be co-regulated with NTRC and THF1 (Figure 3c). THF1 is known to be involved in stabilization and/or assembly of the FTSH subunits in Arabidopsis (Zhang et al., 2009) and Synechocystis (Bec Kova et al., 2017), which are essential for the D1 repair cycle and therefore HL acclimation. In our data, FTSH1 and 2 also cluster together with CRX, albeit FTSH1 is significantly down regulated only in IM_crx_ and not in the knockout mutants. Concerning FTSH2, a similar trend as for CRX is evident. The strongly impaired ability of CRX mutants to grow in HL conditions therefore may also be partly due to a defect in photosystem II repair, at least in IM_crx_ and E1, similar to the THF1 phenotype in Arabidopsis (Sakamoto et al., 2002). Accordingly, PSII proteins were significantly lower abundant in IM_crx_ and E1 than in the respective WTs. However, the A5 mutant did not show a decrease in PSII as compared to CC124 (Figure 3b), thus indicating, that the PSII phenotype does rather dependent on genetic background differences than solely on CRX depletion. Also, other proteins such as RuBisCO and chloroplast ATP synthase (Fig. 3, Supplementary Table 1) were decreased in IM_crx_ and E1 but not in A5, again pointing to differences that manifested in the absence of CRX, as these phenotypes were recovered in the complemented strains, but varied due to the genetic background of these strains. These differences in genetic compositions are consistent with the extensive natural variations found in *Chlamydomonas* (Flowers et al., 2015). Differences in protein expression were also revealed between CC124 and CC125 (Supplementary Figure 2B). In this light CRX is a capacitor that buffers genetic variations, contributing to phenotypic robustness as described for HSP90 (Queitsch et al., 2002).

### CRX inhibits electron transfer between NTRC and PRX1

As NTRC is also a thioredoxin like protein, and as such harbors two pairs of redox active cysteines, we assessed electron transfer between NTRC and CRX as well as PRX1 (Figure 4). Our data revealed that NTRC and CRX did not efficiently exchange electrons. Instead, the presence of CRX inhibited electron transfer from NTRC to PRX1. The degree of inhibition was positively correlated with the amount of CRX added. Considering the amounts of NTRC and CRX as determined by absolute protein quantification in whole cells (Figure 5), NTRC should be mostly inhibited by CRX in HL conditions, assuming the two proteins are in close vicinity to each other inside the chloroplast. The NTRC inhibition by CRX was independent of the CRX redox active site, and therefore cannot be attributed to a direct interaction between the redox active cysteines of NTRC and CRX. Yet, the presence of Ca^2+^ ions had a significant impact on the inhibitory action of CRX towards NTRC (Figure 4). Ca^2+^ is known to activate the CRX redox activity through a conformational change stretching from the Ca^2+^-binding domain to the redox domain of CRX (Charoenwattanasatien et al., 2020). According to the data presented in here, this conformational change could in fact not only lead to CRX activation, but also allow reversible binding of CRX to NTRC (or PRX1) at a site so far unknown, to regulate the NTRC redox activity towards PRX1. A change in Ca^2+^ concentration within the chloroplast stroma would therefore impact the redox signaling network not only at the level of CRX but also indirectly via CRX at the signaling hub NTRC.

Taking advantage of the C136S mutant of NTRC, harboring an inactive NTR domain (Marchetti et al., 2022), allowed us to further dissect the NTRC-CRX-PRX1 interaction (Figure 6), as NADPH-dependent re-reduction of the still active NTRC-TRX domain is disabled in this mutant. We observed reduction of oxidized PRX1 by reduced NTRC and in presence of NADPH as expected, except for the sample containing NADPH and reduced NTRC-C136S. Although reduced NTRC-C136S alone was able to reduce oxidized PRX1, the additional presence of NADPH inhibited this PRX1 reduction (Figure 6A). As suggested by Marchetti et al., the binding of reduced NADPH could lead to a conformational change within the NTRC protein to adopt a ‘closed conformation’ and protect its reducing power from the oxidizing environment. As NADPH cannot be oxidized in NTRC-C136S, this ‘closed conformation’ might be stabilized, so that electron transfer between TRX domain of NTRC-C136S and PRX1 is not possible when NADPH is added. Additionally, it should be noted, that NADPH leads to dimerization, and thereby activation, of NTRC by breaking up protein aggregates formed under oxidizing conditions (Perez-Ruiz and Cejudo, 2009).

Once PRX1 was reduced, NTRC and NTRC-C136S were both able to stabilize it (Figure 6B) and inhibit PRX1 dimerization upon addition of NADPH (Supplementary Figure 2). In the presence of redox-inactive CRX, however, PRX1 dimerization was favored in the presence of NADPH (Figure 6D-F). Upon addition of reduced CRX in the presence of Ca^2+^, PRX1 remained reduced as reported in (Hochmal et al., 2016). The finding that NTRC-C136S cannot stabilize reduced PRX1 in the presence of NADPH and CRX (Figure 6D-F) underpins a direct protein-protein interaction either between NTRC-C136S and CRX, which induces a conformational change that does not further allow to stabilize reduced PRX1, or between CRX and PRX1, which prevents NTRC from acting on PRX1.

These results would indicate a dual role of CRX for PRX1 reduction inside the chloroplast, illustrated schematically in Figure 8: In the presence of Ca^2+^, it might (i) directly reduce oxidized PRX1, thereby getting oxidized itself, and/or (ii) inhibit NTRC driven PRX1 reduction in the presence of NADPH by direct protein-protein interaction (Figure 6D-F). The CRX dependent inhibition of PRX1 reduction by NTRC was also valid at low Ca^2+^ (Figure 6F), albeit at a lower intensity (Figure 5a). Accordingly, inactive CRX would lead to a more oxidized pool of PRX1s, as it cannot reduce PRX1 itself plus prevents reduction by NTRC in the presence of NADPH.

The function of CRX would be particularly important under conditions where cellular Ca^2+^ spikes in response to various biotic and abiotic stresses are evoked (see (Kudla et al., 2018; Pirayesh et al., 2021) for recent reviews on Ca^2+^ signaling in plants), because the physiological Ca^2+^ concentration in the stroma, which is at about 100 – 150 nM Ca^2+^ (Johnson et al. 1995), is lower than the Ca^2+^ concentration needed for half-maximal CRX electron transfer rate (about 280 nM free Ca^2+^) (Hochmal et al. 2016). In the presence of elevated stromal Ca^2+^ concentrations, CRX could be reduced via a NADPH driven NTR like reductase allowing CRX to drive PRX1 reduction (Hochmal et al., 2016). Evidence for CRX driven PRX1 reduction is attained by the fact, that PRX1 levels in the CRX KO mutants are significantly more oxidized in high light as compared to WT (Figure 7).

However, keeping in mind that the redox potentials of CRX (−280 mV) and PRX1 (−310 mV) are similar, CRX reduction via PRX1 should also be possible. The oxidation of PRX1 via CRX in the presence of Ca^2+^ (Figure 6D) could be explained accordingly, also because NTRC-C136S stabilized reduced PRX1 in the absence of CRX (Figure 6B), although its NTR domain is not redox active anymore. As mentioned before, the overall CRX reduction level would also critically depend on the stromal Ca^2+^ concentration.

Under conditions where PRX1 gets quickly oxidized (e.g. stresses like HL, heat, ROS burst), the reduced CRX pool would drive rapid re-reduction of PRX1. In this scenario, CRX could be regarded as a Ca^2+^-regulated redox buffer, important for PRX1 redox status control. At the same time, oxidized CRX could as well drive PRX1 oxidation and/or impact PRX1 reduction via NTRC in the presence of NADPH, shortly after PRX1 has accepted electrons from CRX and the Ca^2+^ spike is over. This scenario, in which CRX is a main player, would therefore be important to return to a steady-state redox equilibrium after transient stress signaling.

Binding of CRX to NTRC or NTRC-C136S in the presence of NADPH (Figure 6D-F) induced a conformational change that impacts the reduction and/or redox state of PRX1. Notably, the catalytically active form of NTRC is the dimer, whose formation is induced by NADPH (Perez-Ruiz and Cejudo, 2009). In this way, function of the NADPH-NTRC complex appears to be under strict control of CRX. The involvement of NADPH in the PRX1 redox tuning via CRX and NTRC allows to interconnect the redox state of the three proteins with active photosynthetic electron transport and metabolism, thereby ensuring an economic use of NADPH for PRX1 reduction. In this light, we propose that the Ca^2+^ dependent redox equilibrium of CRX and PRX1 depends also on the availability of NADPH, where CRX additionally suppresses the reduction of PRX via NTRC.

This situation appears to be different from *Arabidopsis thaliana* where NTRC is the most potent electron donor to reduce the oxidized disulfide form of PRX1 (Pulido et al., 2010). In Chlamydomonas, CRX not only controls electron transfer from NTRC towards PRX1 but can also reduce PRX1 efficiently and is about 10-fold more abundant than NTRC. Thus, we suggest that in the presence of NADPH, CRX is the main electron donor for PRX1 reduction under HL, where CRX is also induced in expression. In darkness or low light, NTRC would be fully functional.

In plants, the absence of NTRC causes growth phenotypes under various abiotic stresses (Serrato et al., 2004; Perez-Ruiz et al., 2006; Chae et al., 2013), particularly under fluctuating light HL regimes (Naranjo et al., 2016). It was also shown that decreased levels of chloroplast 2-Cys PRX suppress the phenotype of the *A. thaliana ntrc* KO, indicating that NTRC is involved in redox balance of chloroplast 2-Cys PRX and optimal photosynthetic performance (Perez-Ruiz et al., 2017). In the absence of CRX, we observed also a more oxidated state of PRX1 (Figure 7), thus suggesting that CRX, in conjunction with NTRC, is likely involved in redox balance of chloroplast PRX1 and the optimal function of the photosynthetic apparatus. This is exemplified in a strong impact in CO_2_ fixation, that is manifested in IM_crx_ as well as in the two *crx* KO strains (Supplementary Figure 1) and the strong HL phenotype observed here (Figure 2). Thus, these data indicate that CRX is, directly and/or indirectly via NTRC, involved in chloroplast redox modulation of TRX targets, besides its redox regulation of PRX1. This impact on chloroplast CO_2_ fixation under HL indicates the unique importance of CRX for the robustness of the system. In line, neither NTRC accumulation nor growth of CRX mutants is affected under low irradiances (Figures 2 + 5). Despite the different protein profiles of the CRISPR/Cas9 and insertional mutant, the strong HL growth phenotype is particularly obvious in the CRISPR/Cas9 mutants and rescued in the A5 complemented strain.

Thus, in conclusion, the KO of CRX is responsible for the HL and CO_2_ fixation phenotypes, possibly linked to elevated ROS formation as described (Hochmal et al., 2016), but not to the PSII decrease in the *crx* KO E1. The impact of the genetic WT background, as already discussed, is striking and indicates that in general observed phenotypes have to be taken with caution, especially when regulatory proteins are knocked-out but also when gene rescue approaches are undertaken. For example, in the IM_crx_, PsbD and PsbC are significantly diminished in HL, yet, rescued in IM_crx_R2. However, PsbD and PsbC are not impacted in expression in the KO A5 under HL (Supplementary Table 1).

In summary, our investigations revealed a novel role of CRX in redox homeostasis, as it impacts NTRC function towards PRX1, adding another layer of complexity to the chloroplast redox and signaling network. This CRX-dependent chloroplast redox regulation is essential for efficient photosynthesis in *Chlamydomonas*.

## Materials and methods

### Strains and culture conditions

The CRX insertional mutant (IM_crx_) with residual CRX accumulation was generated previously (Hochmal et al., 2016) from Chlamydomonas strain CC-4375 (designated as WT_crx_ throughout this manuscript). The mutants E1 and A5, affected in exon 1 and exon 4, respectively, were generated via the CRISPR-Cas9 method from the wild-type strain CC125 (mt+) as described before (Greiner et al., 2017; Kelterborn et al., 2022), to obtain complete *crx* knock-outs (KOs). These strains were then crossed three times with the CC124 (mt−). Progenies without CRX accumulation as evidenced by protein immunoblotting were crossed back into CC125 (mt+), checked again and crossed a third time into CC124 to generate mutants with a clear genetic background. Expression of *crx* was partially restored in the strains IM-R, IM-R2 and A5-R. This was done by introduction of the endogenous *crx* gene (harboring only the first intron) including additionally 900 bp upstream (putative promotor region) and 300 bp downstream (possible additional region important for expression regulation) of the gene to IM_crx_ and A5, respectively. The construct was delivered together with a zeocine resistance cassette to screen for successfully transformed mutants that incorporated the gene into their genome. Induction of expression and expression level was checked by protein immunoblotting and mutants with highest expression level were selected for further analysis.

Strains were grown at 25°C and 120 rpm photo-heterotrophically in tris-acetate-phosphate (TAP) medium or in tris-phosphate (TP) medium to induce photoautotrophic growth. Light intensities were low (25 - 40 μEm^2^s^−1^, LL) or high (200 – 500 μEm^2^s^−1^) as indicated.

### Growth test

Cells were grown under LL in TP medium for three days until a chlorophyll concentration of 3 μg/ml was reached. 10 μl of these cultures were spotted onto two TP or HSM plates each, together with the same volume of culture diluted as indicated. One plate was placed in low light, while the other was incubated under HL (500 μE m^−2^ s^−1^).

### Sample preparation for whole proteome analysis

Cells were grown at 40 μmol photons·m^−2^·s^−1^ (16 h light/8 h dark) for three days in TP and were then adjusted to 4 μg/ml chlorophyll in fresh TP after approx. 4 h in the light cycle. Immediately afterwards, LL samples were harvested and the remaining culture was shifted to HL (500 μEm^−2^s^−1^) for 24 h. Depending on the culture concentration, 10 to 20 ml sample were taken at 0 or 24 h HL by centrifuging at 2500 *x g* for 5 min at room temperature. Pellets were frozen in liquid nitrogen and stored at −80°C until use. Thawed pellets were resuspended in a 1:5 pellet: lysis buffer ratio (150-300 μl, 100 mM Tris-HCl, pH 8, 2% SDS, 10 mM NaF, 10 mM sodium pyrophosphate, 10 mM β-glycerophosphate, 1 mM Na-orthovanadate, 1 mM PMSF, 1 mM benzamidine), heated to 65°C for 20 min while shaking at 1000 rpm, sonicated for 5 min in a sonication bath and centrifuged at 14 000 *x g* for 10 min. The protein concentration of the supernatant was determined by bicinchoninic acid assay (Pierce™ BCA Protein Assay Kit, Thermo Fisher Scientific). 100 μg of protein were loaded into FASP filters (Amicon Ultra 30K, 0.5 ml, Merck) and tryptically digested over night following the FASP (Filter-Aided Sample Preparation) protocol as described in (Wisniewski et al., 2009) with the following modifications: dithiothreitol (DTT) and iodoacetamide (IAA) were replaced by tris(2-carboxyethyl)phosphine) (TCEP) and chloroacetamide (CAA), respectively. allowing reduction and alkylation of cysteines in a single step.

Prior to MS analysis, aliquots corresponding to 5 μg of digested peptides were desalted on C18 membranes as described (Rappsilber et al., 2003), dried by vacuum centrifugation and stored at −80°C until further use. Prior to LC-MS/MS analysis samples were reconstituted in 5 μl of 0.05% trifluoroacetic acid (TFA)/4% acetonitrile (ACN).

### LC-MS/MS analysis of whole proteome

Sample analysis was carried out on an LC-MS system consisting of an Ultimate 3000 nanoLC (Thermo Fisher Scientific) coupled via a Nanospray Flex ion source (Thermo Fisher Scientific) to a Q Exactive Plus mass spectrometer (Thermo Fisher Scientific).

Approximately 1 μg of peptides were loaded on a trap column (C18 PepMap 100, 300 μM × 5 mm, 5 μm particle size, 100 Å pore size; Thermo Fisher Scientific) using loading buffer (0.05% trifluoroacetic acid (TFA)/4% acetonitrile (ACN) in MS grade water) for 3 min at a flow rate of 10 μl/min. Peptides were separated on a C18 column (PepMap 100, 75 μm × 50 cm, 2 μm particle size, 100 Å pore size; Thermo Fisher Scientific) at a flowrate of 250 nl/min. The eluents were 0.1% formic acid (FA) in MS grade water (A) and 0.1% FA/80% ACN in MS grade water (B). The following gradient was applied: 5-24% B over 120 min, 24-36% B over 40 min, 36-99% B over 10 min, 99% B for 20 min. MS data were collected by data-dependent acquisition (DDA), dynamically choosing the 12 most abundant ions from the precursor scans (scan range *m/z* 350–1,400, resolution 70,000, AGC target value 3e6, maximum injection time 50 ms) for fragmentation (MS/MS) by higher-energy C-trap dissociation (27% normalized collision energy, isolation window 1.5 *m/z*, resolution 17,500). AGC target value for MS/MS was 5e4 at 80 ms maximum injection time and an intensity threshold of 6.9e3. Dynamic exclusion was enabled with an exclusion duration of 60 s. Singly charged ions, ions with charge state 5 and above as well as ions with unassigned charge states were excluded from fragmentation. Internal lock mass calibration was enabled on *m/z* 445.120025.MS raw files were converted to mzML format employing ThermoRawFileParser (version 1.3.3, (Hulstaert et al., 2019). Data was analyzed with MSFragger 3.3 (Kong et al., 2017), Philosopher 4.0 (da Veiga Leprevost et al., 2020), and IonQuant 1.7.5 (Yu et al., 2020), all implemented in Fragpipe (version 16.0). The Fragpipe profile “LFQ-MBR” was used with default settings for protein identification and label-free quantification. Spectra were searched against a *Chlamydomonas reinhardtii* protein sequence database consisting of nuclear-encoded (www.phytozome.org, assembly version 5.0, annotation version 5.6), chloroplast-encoded (NCBI BK000554.2), and mitochondria-encoded (NCBI NC_001638.1) proteins. Common contaminants (cRAP, www.thegpm.org/crap/) were also included. Carbamidomethylation was set as fixed modification. Oxidation of methionine and acetylation of protein N-termini were considered as variable modifications.

LFQ data (combined_protein.tsv) was loaded into Perseus 1.6.15.0 (Tyanova and Cox, 2018) for processing and statistical analysis. After log2-transformation, proteins with less than three valid intensity values in at least one strain were filtered out. Then, intensity data from strains with identical genetic background were grouped and proteins not meeting a threshold of 75% of valid values in at least one group were discarded. CRX was exempted from filtering. Remaining missing values were imputed from a normal distribution (width: 0.3, down shift: 2.5). Subsequently, multiple sample testing (ANOVA) was performed with a permutation-controlled FDR of 0.01, followed by Tukey’s honestly significant difference test (FDR 0.05). After z-scoring of protein intensities, hierarchical clustering (average linkage, Euclidean distance, 30 clusters) of significantly differentially expressed proteins was carried out. Volcano plots were generated using LFQ-Analyst (Shah et al., 2020) which uses the Limma (Ritchie et al., 2015) package for differential expression analysis. Imputation of missing data was omitted in LFQ Analyst since this step was already performed in Perseus. False discovery rate (FDR) correction was carried out using the Benjamini-Hochberg method. Cut-offs for log2 fold changes and p-values are given in Supplementary Figure 2. The mass spectrometry proteomics data have been deposited to the ProteomeXchange Consortium via the PRIDE (Perez-Riverol et al., 2019) partner repository with the dataset identifier PXD038009 and 10.6019/PXD038009.

## Measurement of redox activity

Expression and purification of recombinant proteins incl. mutated versions was described earlier (Hochmal et al., 2016; Marchetti et al., 2022). To determine NTRC redox activity, 400 nM NTRC were reduced by 200 μM NADPH at a free calcium concentration of 0 to 2.5 μM for 10 minutes at RT in 30 mM MOPS, 100 mM KCl, pH 7.2 in the presence of different concentrations of CRX WT, its active site mutant (C238S, C242S named C1,2S here), TrxF and TrxR, the *in vitro* electron donor of CRX, as indicated in the corresponding figures. After addition of 200 mM DTNB, the formation of TNB^−^ was measured at 412 nm (Ultrospec 3000, Amersham Biosciences). The reduction rate of DNTB was determined over a time course of 0–80 s after addition of DTNB. The different activities were normalized to the highest activity measured for each batch of purified NTRC. For CRX, 1 μM CRX were reduced as described for NTRC in presence of 200 nM TrxR and different concentration of NTRC-C136S. Reaction and reduction rate were carried out and calculated as described.

### PRX1 interaction assay

The interaction between either NTRC and/or CRX and PRX1 was measured *in vitro* via a photometrical assay (Nelson and Parsonage, 2011). 5 μM recombinant NTRC were reduced at RT by incubation with 120 μM NADPH and 80 μM H_2_O_2_ in 30 mM MOPS, 100 mM KCl, pH 7.2 at a free calcium concentration of 2.5 μM in the presence of different concentrations of CRX WT and TrxR. The NADPH consumption was measured at 340 nm until a steady decrease in absorption was detected. The reaction was then started by addition of 1 μM recombinant PRX1 and recording of the absorption at 340 nm was continued. The rate of NADPH oxidation was calculated from the first 60 s after addition of PRX1.

### NTRC and CRX absolute quantification by Mass Spectrometry

The LC-MS/MS system for sample analysis was identical to the one used for the whole proteome analysis. Recombinant CRX and NTRC (vectors described in (Hochmal et al., 2016) and (Marchetti et al., 2022) were expressed by *E. coli* cultures grown in ^15^N-labelling medium following (Nikolova et al., 2018). Protein purification was done as described ((Hochmal et al., 2016; Marchetti et al., 2022). Cell lysates obtained from four independent unlabelled (^14^N) biological replicates were mixed at equal ratio. Mixed samples were then divided into three aliquots, each containing 50 μg protein (^14^N) plus a known amount (25, 125 or 500 fmol) of the two ^15^N-labelled proteins. Samples were then subjected to tryptic digestion following the FASP protocol as described above. Peptide aliquots corresponding to 5 μg of total protein (^14^N) were desalted, dried by vacuum centrifugation and resuspended in 5 μl of loading buffer (see above). A samples volume of 1 μl was injected, corresponding to 1 μg of ^14^N-labelled peptides plus 0.5, 2.5 or 10 fmol of ^15^N-labelled CRX and NTRC peptides. The mass spectrometer was operated in parallel reaction monitoring (PRM) mode, targeting the light and heavy versions of five distinct peptides per protein. MS raw files were loaded into Skyline (MacLean et al., 2010) for peak extraction and integration of ^14^N and ^15^N peptide signals. From the peptide peak areas of four ^15^N peptides for each protein, calibration curves were generated, which were then used to calculate the absolute amounts of each ^14^N peptide. The protein amount corresponded to the mean value of the peak areas of the respective four ^14^N peptides of CRX and NTRC. Standard deviations relate to the results obtained from the calculations based on the four different peptides used.

### Non-reducing SDS-gels

To visualize the PRX1 redox state, non-reducing SDS-gels were run by using normal Laemmli-loading buffer but omitting DTT and β-mercaptoethanol. 1 μg of recombinant PRX1 (vector and purification described in (Charoenwattanasatien et al., 2018)) was reduced or oxidized with 1 mM DTT or H_2_O_2_ for 1 h at RT. The reducing/oxidizing agent was washed out over a G25 column (GE Healthcare) following the manufacturer’s instructions using 30 mM MOPS, 100 mM KCl, pH 7.2 as buffering agent. Equal amounts of CRX, NTRC or mutated versions were mixed with reduced or oxidized as PRX1 as indicated in the figure. After incubation for 1 h at RT, non-red loading dye was added and samples were heated to 70°C for 20 min before loading onto SDS-PAGE gels.

### NEM-labeling

Recombinant PRX1 was blocked in its respective redox state by TCA precipitation at a final concentration of 10%. After incubation at −20°C for at least 2 h and centrifugation (16 000 × g, 15 min, 4°C), precipitates were washed with 1% TCA in acetone, centrifuged again and then washed with 100% acetone. Washing steps were performed on ice. After another centrifugation step, acetone was removed and the pellet was let dry for approx. 20 min at RT. Pellets were resuspended in 50 mM N-ethyl-maleimide (NEM) in 8 M Urea, 2% SDS, 100 mM Tris/HCl, pH 7.5, 10 mM EDTA for 2 h at RT to block reduced cysteines. After another TCA precipitation step (10% final concentration, 2 h −20°C followed by centrifugation as described above), pellets were washed with 100% acetone twice and resuspended in non-reducing loading dye for SDS-PAGE.

### LC-MS/MS analysis of NEM-labeled PRX1

Gel bands containing PRX1 were cut out of the gel and subjected to tryptic in-gel digestion according to established protocols, and including the reduction and alkylation of non-NEM-labelled cysteines (Shevchenko et al., 2006). After desalting of resulting peptides using C18 STAGE tips (Rappsilber et al., 2007), LC-MS/MS analysis was carried out on the system described above. Samples were loaded on the trap column for 5 min at a flow rate of 10 μl/min using 2.5% loading buffer. Then peptides were separated with the following gradient: 2.5-40 % B over 55 min, 40-99 % B over 5 min, 99 % B for 20 min. MS data acquisition settings were as described for the whole proteome analysis with the following changes: MS/MS maximum injection time was 55 ms and the intensity threshold was 1e4. Dynamic exclusion was set to “auto” assuming a chromatographic peak width (FWHM) of 10 s.

Spectra files were searched in Proteome Discoverer 2.4 against the polypeptide sequence of recombinant PRX1, common contaminants (cRAP, www.thegpm.org/crap/), and the *E. coli* proteome (Uniprot ID UP000000625) using the MSFragger, MS Amanda and Sequest HT nodes. Default search parameters for data acquisition in the Orbitrap were used with the following exceptions, all of which regarding the modification of cysteine residues: NEM-labeling, carbamidomethylation, oxidation, dioxidation and trioxidation were chosen as variable modifications. Peptide precursor intensities were determined via the Minora feature detector node in Proteome Discoverer. Ion traces of the NEM-labeled and the carbamidomethylated versions of the PRX1 peptide VLQAIQYVQSNPDEVCPAGWKPGDK were exported directly from the software. The double peak observed for the NEM derivative is most likely due to the presence of two diastereoisomers resulting from the introduction of a new chiral centre by the labelling (Jemal and Hawthorne, 1994; Srinivas and Mamidi, 2003).

### MIMS

To compare the physiology of the mutated CRX strains (E1 and A5), to their background wild-type strain (CC125) as well as IM_*crx*_ and IM-R2, cells were grown in TP medium and kept at 24.5°C under a continuous irradiance of 30 μmol photons·m^−2^·s^−1^ (referred to as low light, LL). When stated, cells were exposed to 500 μmol photons·m^−2^·s^−1^ for 12 h prior to measurements (referred to as high light, HL). The cells were then harvested, centrifuged at 3300 × g for 2 min and resuspended to a final concentration of 15 μg/ml chlorophyll in TP medium, supplemented with 50 mM HEPES and 2 mM Na_2_CO_3_, at pH 7.2. Photosynthetic activity was assessed by a combined home-built membrane inlet mass spectrometer (MIMS) and a dual pulse amplitude modulated fluorometer (PAM). The measured masses were, 32, 36, 40, and 44, correlating to ^16^O_2_, ^18^O_2_, Ar and CO_2_ respectively. The cells were then maintained in a short dark regime, during which their quantum efficiency (F_v_/F_m_) was determied by exposing them to a saturating pulse (10 mmol photons·m^−2^·s^−1^, 300 ms). During measurements, the cells were exposed to high light (500 μmol photons·m^−2^·s^−1^, mixed red/blue light) for 3 min, followed by darkness for 3 min. This scheme was repeated twice, in order to decrease noise and verify that no further changes were generated by the short light exposure time. Net O_2_ evolution was assessed by tracking ^16^O_2_ concentration during light exposure. Gross O_2_ evolution was assessed by subtracting ^18^O_2_ uptake in light relative to dark exposures, from net O_2_ evolution, as was previously demonstrated in (Kedem et al., 2021). CO_2_ fixation was assessed by subtracting the rate of CO_2_ uptake during light exposure, from its evolution rate under darkness.

## Author contributions

M.H., K.Z., P.H. designed the experiments; K.Z., G.M.M., R.F., Y.M. performed the experiments; A.O., S.K. generated the CRISPR/Cas9 knockout strains, M.H., K.Z., G.M.M., Y.M., M.S. analyzed the data; K.Z., G.M.M. visualized the data; K.Z., M.H. wrote the manuscript with contribution of all authors; I.Y., P.H., M.H. acquired the funding.

---

**Supplementary Figure 1: Decreased photosynthetic activity of *crx* mutants**. CRISPR-Cas9 *crx* mutants A5 and E1, and the parental strain CC125 as well as WT_crx_, IM_crx_ and complemented strain IM-R2 were exposed to light following 12 hours of low (30 μmol photons·m^−2^·s^−1^, LL) or high (500 μmol photons·m^−2^·s^−1^, HL) exposure. Quantum efficiency was determined by exposing the cells to a saturating pulse previous to illumination (A). Simultaneously, dissolved O_2_ and CO_2_ concentrations were evaluated by MIMS. O_2_ net evolution rate was determined by the increase of ^16^O_2_ concentration during light exposure (B). The gross O_2_ production rate was determined by subtracting the rate of ^18^O_2_ decrease in light (termed as ‘total uptake’), from the net O_2_ evolution (C). CO_2_ fixation was determined by subtracting the rate of CO_2_ uptake in light from its increase under darkness (D). Each column represents the average of at least 5 biological replicates, statistical significance was determined using a student’s T-test, with asterisks indicating a p-value of *<0.05 and ***<0.001.

**Supplementary Figure 2: Volcano plots of quantitative (LFQ) proteomics data**. Log2 fold changes in protein abundances are plotted against Benjamini-Hochberg-adjusted p-values. Panel **(A)** displays combined data of all mutant strains versus all wildtypes. NTRC1 and FTSH1 (Cre01.g066552) are significantly less abundant in *crx* mutants (adj. p-value <0.01; FC_*crx*/WT_ <−0.5, >0.5; n=12). **(B)** Volcano plot to identify differences in the proteomes of CC124 and CC125 wildtype strains. Proteins involved in acetate metabolism/glyoxylate cycle (red rectangles) are much more abundant in CC124. Differentially expressed proteins (adj. p-value <0.01; FC _CC124/CC125_ <−1, >1; n=4) are highlighted in black.

